# Development of a novel rodent rapid serial visual presentation task reveals dissociable effects of stimulant vs non-stimulant treatments on attention

**DOI:** 10.1101/2021.10.08.463723

**Authors:** Abigail Benn, Emma S.J. Robinson

**Affiliations:** University of Oxford, Department of Experimental Psychology, Tinsley Building, Marsden Road, Oxford, OX1 3TA; University of Bristol, School of Physiology, Pharmacology and Neuroscience, Biomedical Sciences Building, University Walk, Bristol, BS8 1TD

**Keywords:** RSVP, attention, amphetamine, atomoxetine, CPT

## Abstract

The rapid serial visual presentation (RSVP) task and continuous performance tasks (CPT) are used to assess attentional impairments in patients with psychiatric and neurological conditions. This study developed a novel touchscreen task for rats based on the structure of a human RSVP task and used pharmacological manipulations to investigate their effects on different performance measures. Normal animals were trained to respond to a target image and withhold responding to distractor images presented within a continuous sequence. In a second version of the task a false-alarm image was included so performance could be assessed relative to two types of non-target distractors. The effects of acute administration of the stimulant and non-stimulant treatments for ADHD (amphetamine and atomoxetine) were tested in both tasks. Methylphenidate, ketamine and nicotine were tested in the first task only. Amphetamine made animals more impulsive and decreased overall accuracy but increased accuracy when the target was presented early in the image sequence. Atomoxetine improved accuracy overall with a specific reduction in false-alarm responses and a shift in the attentional curve reflecting improved accuracy for targets later in the image sequence. However, atomoxetine also slowed responding and increased omissions. Ketamine, nicotine and methylphenidate had no specific effects at the doses tested. These results suggest that stimulant versus non-stimulant treatments have different effects on attention and impulsive behaviour in this rat version of an RSVP task. These results also suggest that RSVP-like tasks have the potential to be used to study attention in rodents.

## Introduction

Attentional impairments are observed across a wide range of psychiatric and neurological disorders. The rapid serial visual presentation task (RSVP) and continuous performance tasks (CPT) have been used to study neuromodulatory systems implicated in attention [1–3], or disorders involving perturbed attentional processing [4–7]. In these tasks inattention reflects a failure to respond to a target stimulus (errors of omission) and inhibit responding to non-targets (errors of commission) [4, 8]. Drugs modulating attention have been shown to affect one or both of these performance measures [9–11].

There have been a number of different attentional tasks developed for rodents [8, 12–14], and one of the most widely used is the 5 choice serial reaction time task (5CSRTT) which measures visuo-spatial attention in rodent requiring attention to a light cue presented in one of five spaced apertures [15]. However, several challenges still exist with, these tasks including the use of discrete trials, fixed intervals and extensive training methods which can lead to procedural learning and timing strategies [16] although use of a variable inter-trial interval can help to mitigate some of these issues [17, 18]. In order to look at ways to increase the attentional demands of the 5CSRTT and reduce animals reliance on a timing strategy, a 5-choice continuous performance task (5CCPT) has been developed which includes no-go (non-target) trials to align more with response indices in human tasks e.g. Connor’s CPT [8, 11, 19]. There has also been rodent touchscreen CPT (rCPT) developed which consists of presenting several images (target or distractor images) across a single trial but with an ITI between image presentations [13] or a target image with flanking distractors [14]. Performance parameters such as false-hit and correct-rejections are used to infer discrimination sensitivity (d’) from Signal Detection Theory, made possible by presentation of discrete go (target) and no-go (distractor) trials [13, 20–22]. Similar to a concept of a touchscreen CPT, we hypothesized that training animals in a task using a randomized image sequence (RSVP stream), containing a target and multiple distractor images would result in a challenging and potentially translational rodent equivalent of this human attentional task. In our task we chose stimuli presentation that resembles the single-target human RSVP attention task, where participants respond to an unpredictable target image embedded within distractor sequences, for origins of the task see [23–25]. Hit rate (accuracy) and false-alarm rate are used to quantify the subject’s target detection performance within the RSVP stream [26]. The position of the target image is temporally distributed within the image sequence and pseudo-random in relation to the positions of the distractor images [4].

We designed a touchscreen-based rodent task using a continuous sequence of images where rats were trained to recognize a single image as the target whilst withholding from responding to the other images. Our prediction was that this task would be less reliant on timing strategies, require animals to sustain attention for a longer period of time, and might enable us to dissociate between mechanisms involved in enabling animals to sustained attention for the duration of the sequence (attention curves plotting accuracy against time), discriminative or perceptual accuracy (target versus false alarm image), and impulsive responding (where responses increase irrespective of target category). To test this, we first trained animals using a random sequence of images containing a single target and five non-target images. Based on these initial findings, we added one distractor with an image which contained many of the same features as the target to provide a false alarm image. We hypothesized that this would give a clearer distinction between target (accuracy), false-alarm, and distractor responses, and the ability to distinguish inattention (responses to false-alarm image) versus impulsive responding (responses to all distractor images). We tested different pharmacological treatments which have been studied in similar rodent attention tasks and with relevance to human ADHD [3, 27–29] and to compare the effects of stimulant versus on-stimulant ADHD medications. In should be noted that a limitation of the current study was that only atomoxetine and amphetamine were tested in both cohorts. The design of the task offers both advantages and disadvantages relative to other rodent attention tasks and is not meant to be a replacement but rather add an additional option for study different aspects of attentional processing in rodent models. In this initial study only male rats of a single strain were tested and further studies in different strains, sexes and also mice are necessary to establish greater validation.

## Methods

### Subjects

Subjects, housing and husbandry were similar to previous published studies^14^. Two cohorts of male Lister hooded rats (*n*=12 per group) weighing approximately 300g at the start of training (Harlan, UK). Rats were pair housed with standard environmental enrichment (bedding, cardboard tubes) under temperature-controlled conditions and 12-hh reverse light-dark cycle (lights off at 0700h). Rats were food restricted to approximately 90% of their free feeding weight (∼18g/day laboratory chow, Purina, UK), with water provided ad libitum. Procedures were conducted and are reported in accordance with the ARRIVE guidelines and requirements of the UK Animals (Scientific Procedures) Act 1986, and approved by the University of Bristol Animal Welfare and Ethical Review Board. Behavioral testing was carried out between 8am and 5pm during the animals’ active phase.

### Behavioral Training

A rat-rapid serial visual presentation task (R-RSVP) was designed based on the human RSVP task [62] and used a single target [4] and therefore differs from the attentional blink paradigm [1, 56].

Touchscreen boxes (Med Associates, USA) containing three screen panels (left, right, center), controlled by KLimbic Software (Conclusive Solutions Ltd, UK) were used for all training and testing. The behavioural equipment and software were supplied by OCB Solutions Ltd, European distributors for Med Associates UK (https://www.med-associates.com/contact/). Rats were trained to screen press in response to a specific stimulus image (‘spider’) embedded in a sequence of distractor images (Fig 1). The training schedule used for both cohorts is described in detail under supplementary methods and summarized in supplementary table S1. All rats from cohort 1 and 2 were successfully trained using this graduated training procedure over a period of 42-60 sessions. Rats were trained using images (jpeg 260×380 pixels) illustrated in fig 1a (cohort 1) and fig 1b (cohort 2). Each cohort responded to the same target image (‘spider’). The introduction of an image that more closely resembled the target image versus the other distractor images was implemented to act as a false alarm (see video 1). This was designed to resemble instances where certain targets and distractors are closely related, for example letter ‘S’ and numeral ‘5’ [63] making accuracy identification of the target harder. All animals were trained using the same image set and target versus non-target images were not counter-balanced. There is the potential for perceptual differences to impact on the performance and the ease with which animals differentiate between images and this can be seen with cohort 1 where the distractor images are not achieving the same level of performance (Fig 2). For cohort 2, we changed some of the images and also added a false alarm and this achieved a baseline performance more in line with what we would predict based on performance in humans. Because we have used 6 different images, and the false-alarm image is paired with the specific target, to fully counter-balance design would require a very large number of combinations. The design used here may introduce a bias related to the specific choice of target and non-target images however, we use a within-subject design which will help to reduce potential confounds this could introduce although not completely mitigate these. Future studies could look in more detail at the images being used and optimize for perceptual similarity and a design which is more readily counter-balanced.

**Figure 1.**
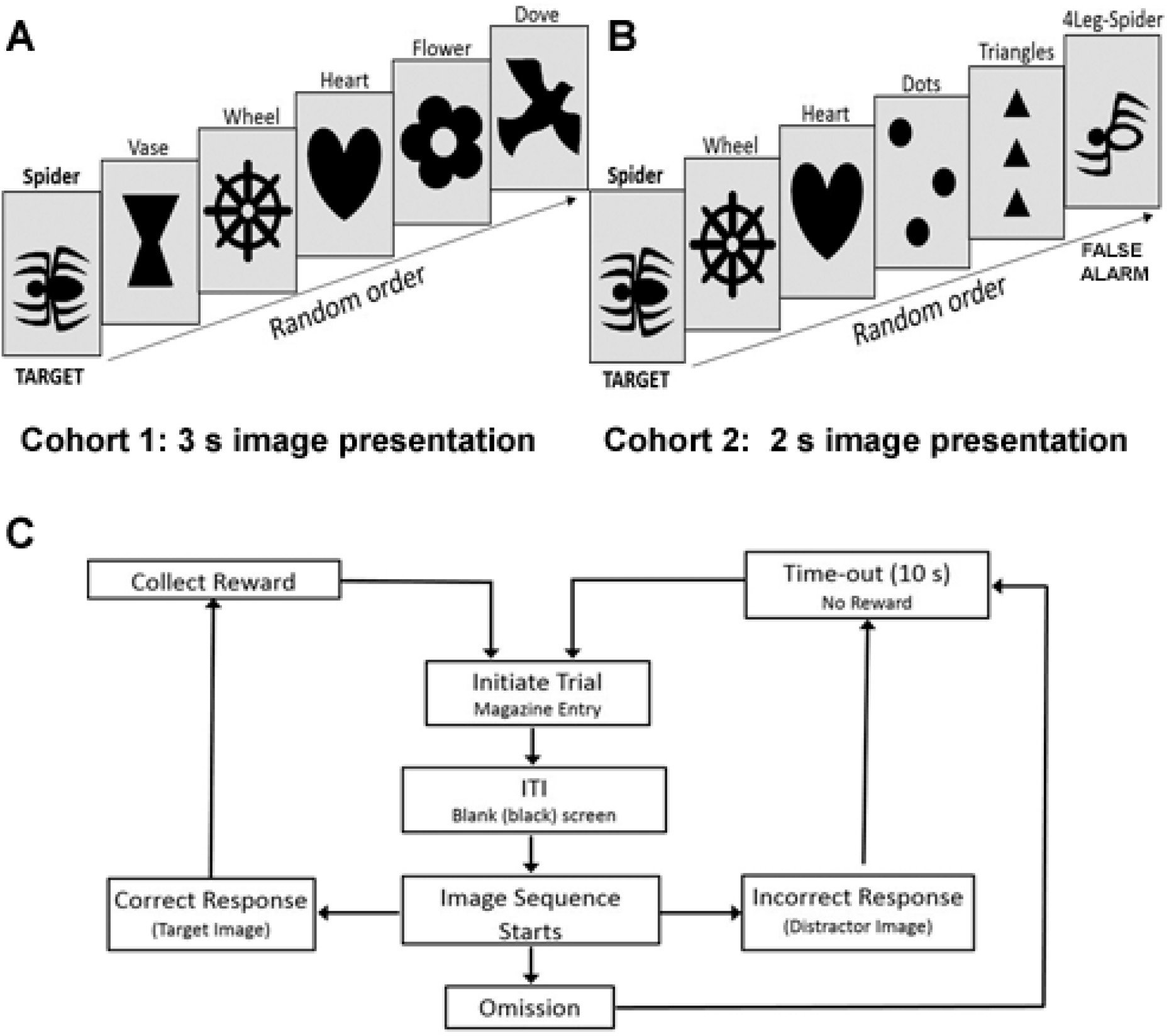
Images and trial outcomes for the rat-rapid serial visual presentation task (R-RSVP). Images used for cohort 1 (a) and cohort 2 (b) with presentation time of 3 s and 2 s respectively. Cohort 2 images contained a false-alarm (4-leg spider) image. Flow chart representing all possible trial outcomes during task performance (c).

**Figure 2.**
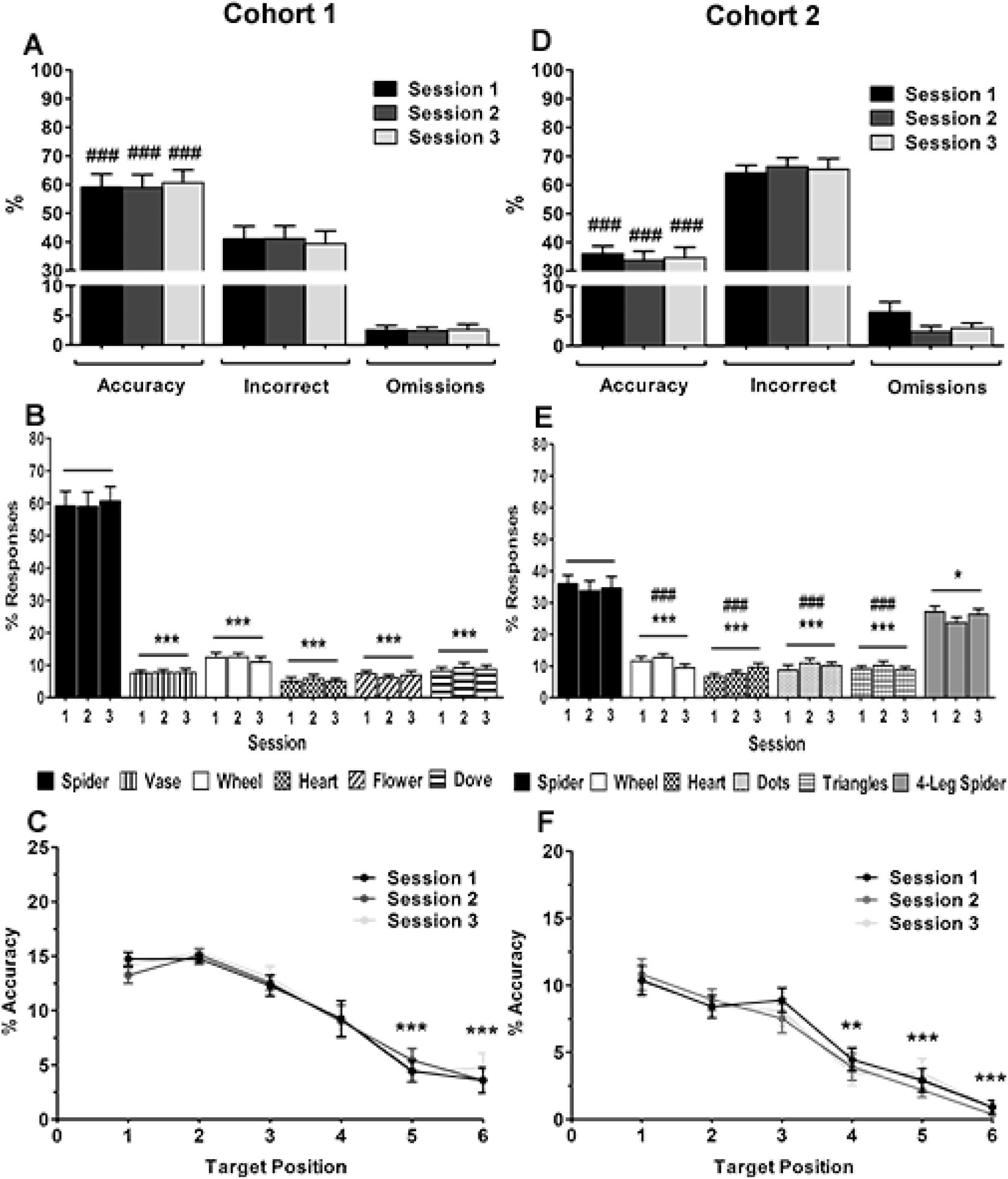
Performance data for cohort 1 (a-c) and cohort 2 (d-f) for the last three consecutive pre-drug baseline sessions. Response data for % accuracy, % incorrect, and % omissions for cohort 1 (a) and cohort 2 (d). Image responses for cohort 1 (b) and cohort 2 (e), spider is the target image (accuracy). The sum of distractor responses (all images except spider) is equivalent to incorrect responses in (a). Attention curves showing accuracy per target sequence position for cohort 1 (c) and cohort 2 (f). Results are shown for the total population, mean ± SEM, *n* = 12 animals per cohort, \**p*<0.05, \*\**p*<0.01, \*\*\**p*<0.001, versus target image (spider) or target sequence positon (within-subject), ^###^*p*<0.001 versus 4-leg spider (false alarm) or chance performance (17%, 1-sample *t*-test).

**Table 1:** Latency data for amphetamine and atomoxetine

| Cohort | Drug | Dose (mg/kg) | Correct Latency (s) | Incorrect Latency (s) | False-Alarm Latency (s) | Response Latency (s) | Collection Latency (s) |
| --- | --- | --- | --- | --- | --- | --- | --- |
| 1 | AMP | 0.0 | 1.10 ± 0.04 | 1.46 ± 0.06 | - | 6.77 ± 0.26 | 1.56 ± 0.09 |
|  |  | 0.3 | 1.04 ± 0.04 | 1.50 ± 0.04 | - | **5.60 ± 0.32\*\*\*** | 1.47 ± 0.08 |
|  |  | 1.0 | 1.09 ± 0.04 | 1.54 ± 0.06 | - | **3.90 ± 0.33\*\*\*** | 1.40 ± 0.10 |
|  | ATO | 0.0 | 0.97 ± 0.06 | 1.53 ± 0.05 | - | 6.13 ± 0.35 | 1.56 ± 0.07 |
|  |  | 0.3 | 1.04 ± 0.06 | 1.49 ± 0.05 | - | **7.18 ± 0.35\*\*** | 1.61 ± 0.06 |
|  |  | 1.0 | 1.07 ± 0.06 | 1.56 ± 0.06 | - | **8.08 ± 0.20\*\*\*** | **1.76 ± 0.07\*\*** |
|  |  | 3.0 | **1.24 ± 0.10\*** | 1.59 ± 0.09 | - | **9.11 ± 0.29\*\*\*** | **1.87 ± 0.07\*\*** |
| 2 | AMP | 0.0 | 0.87 ± 0.05 | 0.98 ± 0.03 | 1.02 ± 0.05 | 3.77 ± 0.22 | 1.62 ± 0.11 |
|  |  | 0.3 | 0.90 ± 0.05 | 1.01 ± 0.04 | 1.00 ± 0.06 | **3.31 ± 0.25\*** | 1.47 ± 0.06 |
|  |  | 1.0 | 0.83 ± 0.04 | 1.01 ± 0.03 | **0.84 ± 0.04\*\*\*** | **2.15 ± 0.18\*\*\*** | 1.46 ± 0.08 |
|  | ATO | 0.0 | 0.92 ± 0.05 | 1.01 ± 0.04 | 1.04 ± 0.04 | 3.98 ± 0.27 | 1.61 ± 0.12 |
|  |  | 0.3 | 0.97 ± 0.04 | 0.96 ± 0.04 | 1.16 ± 0.06 | **4.92 ± 0.26\*** | **1.85 ± 0.12\*** |
|  |  | 1.0 | 0.90 ± 0.05 | 0.92 ± 0.07 | 1.14 ± 0.05 | **4.96 ± 0.30\*\*\*** | **1.83 ± 0.09\*** |
|  |  | 3.0 | 1.00 ± 0.06 | 0.92 ± 0.06 | 1.17 ± 0.11 | **5.59 ± 0.25\*\*** | **1.93 ± 0.11\*** |

The effect of amphetamine (AMP) and atomoxetine (ATO) on latency measures in the rat-rapid serial visual presentation task (R-RSVP) for cohort 1 (3 s image presentation) and cohort 2 (2 s image presentation). Only images used with cohort 2 contained a false-alarm image (4-leg spider) therefore false alarm latency for cohort 1 was not recorded. Results are shown for the total population, mean ± SEM, $n = 12$ animals per cohort, \* $p < 0.05$ , \*\* $p < 0.01$ , \*\*\* $p < 0.001$ , versus vehicle (within-subject).
AMP amphetamine, ATO atomoxetine

### Drugs

Cohort 1 received treatments in the following order: amphetamine, atomoxetine, methylphenidate, ketamine, nicotine. Cohort 2 received treatments in the following order: amphetamine, atomoxetine. Atomoxetine hydrochloride (0.3-3.0mg/kg, t=-40 min), ketamine hydrochloride (1.0-10.0mg/kg, t=-5 min), and nicotine (0.01-0.1mg/kg, t=-10 min) were purchased from Tocris Bioscience, UK, dissolved in 0.9% saline and administered by intraperitoneal injection. Methylphenidate (1.0-10.0mg/kg, t=-30 min) and amphetamine (0.3-1.0mg/kg, t=-30 min) were purchased from Sigma-Aldrich, UK, dissolved in distilled water and mixed with strawberry milkshake (50:50, Yazoo, Campina, UK) for oral administration. All drugs were prepared fresh each day and dosed in a final volume of 1ml/kg. Doses used were based on previous behavioral studies [18, 61, 64, 65] and administered using a refined injection method (Stuart et al., 2015). The choice of doses and route of administration were based on previous studies in the 5CSRTT and evidence that the neurochemical effects of the psychostimulants can be influenced by the route of administration. Specifically, Berridge et al (2006) found that oral administration of methylphenidate resulted in less effects on subcortical dopamine and a relatively selective increase in noradrenaline and dopamine in the prefrontal regions [43]. Whilst the other drugs tested have most commonly been administered by the intraperitoneal route, this does not preclude a possible effect of route of administration for any of the treatments and may lead to some differences in effects in the task which have yet to be explored.

### Testing Procedure

Cohort 1 animals were trained using images shown in fig 1a (3s image presentation) and performed the following dose-response experiments; amphetamine, atomoxetine, ketamine, nicotine, and methylphenidate. Cohort 2 were then trained using images in fig 1b (2s image presentation) and performed amphetamine and atomoxetine dose-response experiments only. Animals received drug doses according to a fully randomized design, with the experimenter blind to treatment. It is possible that animals performance was influenced by the prior drug tested which could have been mitigated if all drugs and doses had been tested in a fully randomized design however, this would have increased the number of factors and reduced power and would have required a much larger sample size. A drug-free baseline session preceded each drug day and a washout day (no testing or drugs) proceeded each drug day. At least 8 days drug-free baseline sessions were performed before commencing the next drug and animal baseline performance analyzed to check they were stable. A RM ANOVA with TIME as factor was used to assess for effects on baseline performance but there were no significant effects observed (data not shown). An acclimatizing dose of nicotine 0.1mg/kg was administered to all rats two days prior to the start of the dosing regimen. The highest dose of nicotine (0.3mg/kg) was administered separately after the lower doses were found to be ineffective.

### Performance Measures

In this new task, we were able to record a number of different parameters which we suggest may align to different aspects of attentional processing and impulsive behaviour. For each trial sequence, a single outcome was recorded and any response terminated the trial sequence with a new sequence initiated by the animal following either consumption of the reward (correct trial) or a time-out (incorrect or omission). Correct or incorrect responses recorded during the sequence also generated a response latency. Omissions were recorded when the sequence reached the end, and no response was recorded. Omission can arise either because the animals fails to detect and respond to the target image or because they are not attending to the screen. Because animals must initiate the start of each sequence of images, total omissions in this task were less influenced by overall task engagement and we can use total trials initiated to make inference about this. When animals make a correct response, latency to collect the reward is also recorded.

Using the data obtained for correct versus incorrect responses we were able to calculate a number of different variables and using this range of measures, can make some inference about the different aspects of attention and impulse control which might influence performance. % accuracy (correct trials divided by total number of correct and incorrect trials *100), % incorrect (incorrect trials divided by total number of accuracy and incorrect trials *100), and % omissions (omitted trials divided by total number of omitted, accuracy, incorrect trials *100) were calculated for the whole session alongside total trials completed similar to the data report for the 5CSRTT and 5-CPT. The normalization of the data takes into account the relative proportion of trials completed and, for correct and incorrect responses, that the same motor effort needed [41]. Data for the individual image responses were expressed as % responses for each image (number of responses per image divided by the total number of image responses for accuracy and incorrect trials*100), where % responses for the target is equivalent to % accuracy, and the sum of the distractor responses is equivalent to % incorrect.

Correct latency and collection latency represent the time taken to respond to the target image and to collect reward respectively, averaged across the total number of accuracy trials. Response latency is the time taken to respond to an image following sequence initiation, averaged across the number of accuracy and incorrect trials. Incorrect latency represents the time taken to respond to a distractor image, averaged across the number of incorrect trials. For cohort 2, the latency to respond to the false-alarm image was calculated separately from the other distractor images (incorrect latency). All latency data are presented in seconds (s) and only those with >0.2s included in analysis, based on minimum time needed to process and initiate a motor response to touchscreen images in rats [40, 66]. Latencies <0.2s were therefore interpreted as delayed responses to the previous image rather than the current image.

Due to the absence of discrete no-go trials, false-hit and correct rejection responses, d’ could not be analyzed compared to other rodent CPTs [13, 20]. However, we were able to plot the data for correct responses relative to the position of the target image in the sequence to obtain attention curves. Attention curves were only calculated where a main effect on accuracy was observed and data were expressed as % accuracy of responding to the target image in each sequence position (1^st^ to 6^th^ image presented). This analysis specifically looked at responses to target images and the resulting attention curve could be influenced by either a change in impulse control, or impairments in sustained attention. However, by also looking at the results for accuracy by image type we, can see whether the pharmacological treatment causes a change in impulsive responding which parallel the effects on attention i.e. are related, or, whether changes in the attention curve arise in the absence of any changes in impulse control. Sustained attention tasks asses the ability of the animal to monitor intermittent and unpredictable events over a sustained period of time (Wicks et al., 2017). By presenting animals with a continuous sequence of random stimuli over a prolonged period but requiring them to monitor and detect the correct stimuli, we suggest that this RSVP task and particularly the attention curves, may provide a measure of sustained attention. Rodent and human psychomotor vigilance tasks (PVTs) require subjects to respond to stimuli randomly presented within a fixed period of time. Decreases in vigilance may be observed by slower reaction times, increases in omissions and increases in premature responding [67]. As for other rodent attention task, in this RSVP task we are able to extract these same measures thus providing us with a measure of vigilance. In the final modification to the task where we introduce the false alarm, we can then measure responses to the target versus this near target image against the other distractor images. This has the potential to provide a measure of discriminative or perceptual accuracy.

### Statistical analysis

Data was formatted and performance measures calculated using MATLAB® for Windows (MathWorks Inc version R2015a, USA, https://uk.mathworks.com/). Pre-drug and dose-response performance measures were analyzed using separate repeated-measures ANOVA (RM-ANOVA) with session or dose as within-subject (ws) factors. A one-sample t-test was used to check performance levels were above chance (17%) during pre-drug baseline performance. Each drug study was analyzed as an independent experiment. Image responses were analyzed using separate RM-ANOVA with session and image, or dose and image, for pre-drug and dose-response data respectively. Accuracy at each target position in the sequence (1^st^ - 6^th^) was analyzed using separate RM-ANOVA with session and position, or dose and position, for pre-drug and dose-response data respectively. Image responses and accuracy curves were analyzed in instances where significant main effects on attention (% accuracy) were found (amphetamine and atomoxetine only).

Data were checked for normality using Shapiro-Wilk and only the data for the attention curve for the high dose of amphetamine in cohort 2 was found to deviate significantly and this was due to one animal which would meet criteria for an outlier (2 standard deviations from the mean). If this animal was removed, the data were normally distributed. As the majority of the data met the requirements for parametric analysis and 2 factor non-parametric ANOVA is not straight forward, we proceeded with RM-ANOVA analysis. Where significant main effects were observed, Sidak post-hoc tests were used to further analyze the differences between groups. Degrees of freedom were adjusted to more conservative values using the Huynh-Feldt epsilon correction for instances of sphericity violation according to Mauchly’s test and the data were checked for the assumption of homogeneity of sample variance using Levene’s test. Epsilon values (ɛ) are stated where the degrees of freedom have been corrected, alpha level was set at 0.05. SPSS for Windows (IBM version 23, USA, https://www.ibm.com/analytics/spss-statistics-software) was used for statistical analysis. Sample size was based on previous studies using similar behavioral tasks. Graphs were plotted using Prism 7 (GraphPad, USA, https://www.ibm.com/analytics/spss-statistics-software).

## Results

### Baseline data

*Cohort 1:* Animals for both cohorts were able to discriminate between the target image and distractors (including the false alarm image used for Cohort 2) and achieved a stable baseline performance (Cohort 1: Fig 2a, Supplementary table S2; F_2,22_ < 0.71, *p* > 0.501; Cohort 2: Fig 2d, supplementary table S2; Session F_2, 22_< 2.93, p > 0.074). Animals could discriminate the target from distractor images above the level of chance (17%) (Cohort 1: Fig 2a; Session 1-3 *t*_11_>9.20, *p*<0.001; Cohort 2: Fig 2d; Session 1-3 *t*_11_>4.68, *p*<0.001). Target responses were significantly higher than responses to distractor images and the introduction of the false alarm with cohort 2 resulted in a higher level of responding for this image relative to the other distractors (Cohort 1: Fig 2b; Image F_1.2, 13.1_ = 89.36, *p* < 0.001, ε = 0.24, Session F_2, 22_ = 3.48, *p =* 0.049, Image*Session F_6.9, 76.3_ = 0.68, *p =* 0.689, ε = 0.69, spider vs vase/wheel/heart/flower/dove *p <* 0.001; Cohort 2: Fig 2e; Image F_1.6,18.1_=38.59, *p* < 0.001, spider vs 4-leg spider *p* = 0.014, 4-leg spider vs all other distractor images *p*<0.001). Using an analysis of accuracy over time, we observed that responses declined with lower accuracy when the image was presented in position 5 or 6 (Cohort 1: Fig 2c; Position F_2.4,26.1_=57.17, *p*<0.001, ε = 0.48, 1 versus 5,6 *p*<0.001, 1 versus 2-4 *p >* 0.05, Session F_2,22_ = 0.71, *p =* 0.500, Session*Position F_10,110_ = 1.23, *p* = 0.278; Cohort 2: Fig 2f; Position F_3.8,41.7_=42.86, *p*<0.001, ε = 0.76, 1 versus 4-6 *p*<0.01, 1 versus 2,3 *p* > 0.05, Session F_2,22_ = 1.07, *p* = 0.359, Position*Session F_10,110_ = 0.70, *p* = 0.720). Baseline data were also analysed for each drug study to check if animal’s performance remained stable across the two-week testing protocol. No significant differences were observed during any of the drug testing protocols (data not shown).

### Amphetamine Dose Response

Similar findings for amphetamine were observed for both cohort 1 and 2. Analyzing the attention curves revealed higher accuracy when the target image was presented in the first position in the sequence but reduced accuracy for the later images. Cohort 2 did not show any relative difference in response errors for false alarm image versus the other distractor.

#### Cohort 1

Amphetamine reduced overall accuracy and increased incorrect responding (Fig 3a; Dose F_2, 22_=64.45, *p*<0.001, vehicle versus 0.3mg/kg *p* = 0.002, 1.0 mg/kg *p* < 0.001), but also reduced omissions (F_1.1,12.4_=8.62, *p*=0.010, vehicle versus 0.3mg/kg *p* = 0.038, 1.0 mg/kg *p* = 0.006). The number of correct trials declined with dose (Supplementary table S3; F_2, 22_=55.72, *p*<0.001, vehicle versus 0.3mg/kg *p* = 0.009, 1.0 mg/kg *p* < 0.001) but the total number of trials animals performed was unaffected by treatment (Supplementary table S3; F_2, 22_=1.94, *p*=0.167). Response latency was reduced at all doses tested (Table 1; Dose F_2,22_=57.71, *p*<0.001, vehicle versus 0.3 mg/kg, 1.0 mg/kg *p* < 0.001) but other latency measures were unaffected (Table 1; Dose F_2, 22_ < 2.86, *p* > 0.079). Amphetamine treatment differentially effected responding depending on the image (Fig 3b; Image F_1.4, 15.3_ = 65.53, *p* < 0.001, ε = 0.28, Dose F_2, 22_ = 3.92, *p* = 0.035, Image*Dose F_10, 110_ = 39.12, *p* < 0.001, ε = 0.68) with increased responding for all but the vase image although this was close to significance (*p*>0.066). Accuracy increased following 0.3 mg/kg dose when the target image was presented at the earliest sequence position (Fig 3c; Dose F_2,22_ = 64.52, *p* < 0.001, Position F_2.8,31.2_ = 148.83, *p* < 0.001, ε = 0.70, Position*Dose F_7.4,81.8_ = 9.59, *p* < 0.001, ε = 0.74, Position 1 *p* = 0.019), but reduced for later target positions (Positions 3-6 *p* < 0.041). The highest dose reduced accuracy when the target image was presented following target positions 2-6 (*p* < 0.001).

**Figure 3.**
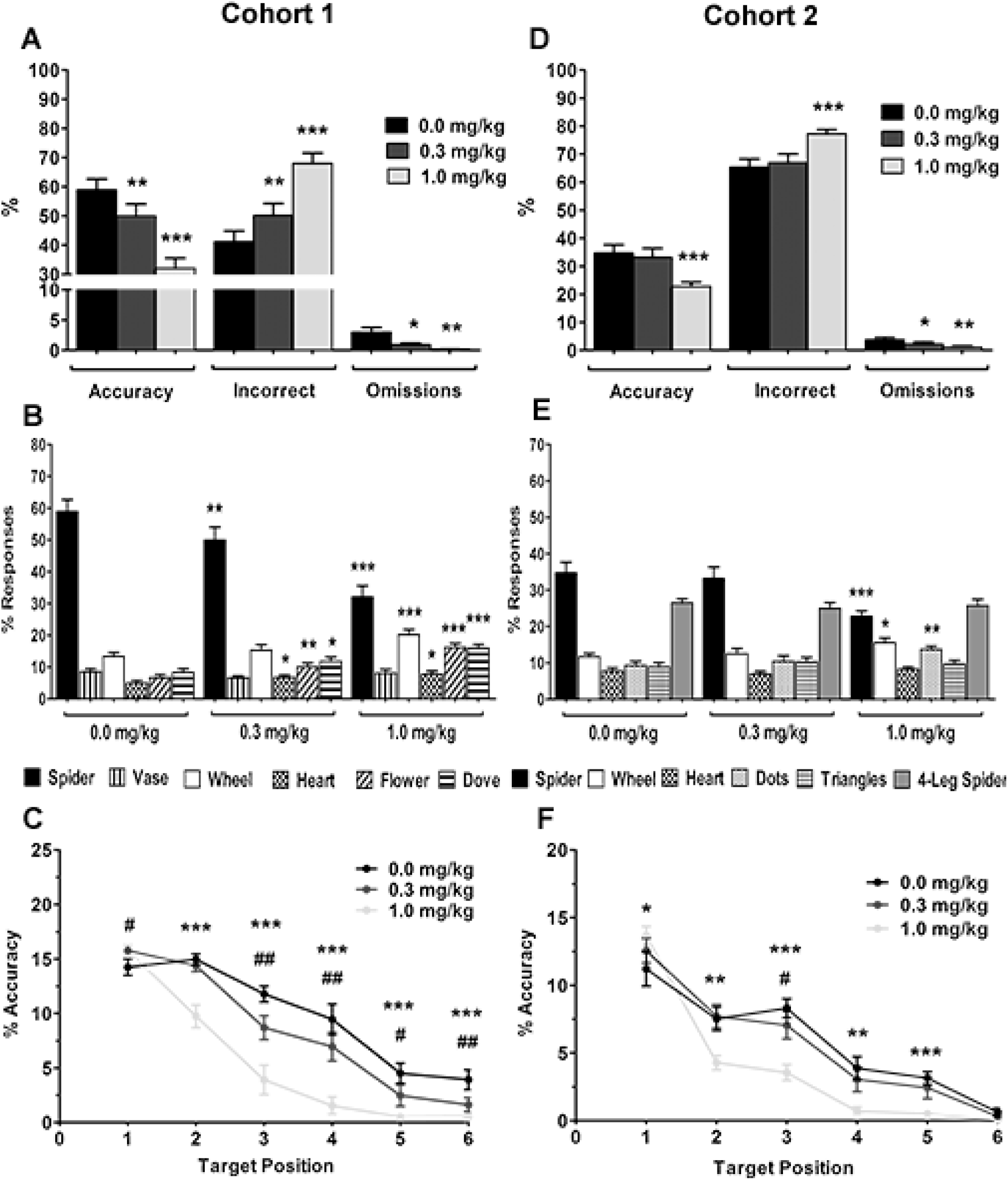
The effect of amphetamine on performance in the rat-rapid serial visual presentation task (R-RSVP). Performance data for cohort 1 (a-c) and cohort 2 (d-f), response data for % accuracy, % incorrect, and % omissions for cohort 1 (a) and cohort 2 (d). Image responses for cohort 1(b) and cohort 2 (e), spider is the target image. The sum of distractor responses (all images except spider) is equivalent to incorrect responses for each dose in (a). Attention curves showing accuracy per target sequence position for cohort 1 (c) and cohort 2 (f). Results are shown for the total population, mean ± SEM, *n* = 12 animals per cohort. Response data (a,b,d,e); \**p*<0.05, \*\**p*<0.01, \*\*\**p*<0.001, versus vehicle (within-subject). Accuracy per target sequence position (c,f); #*p*<0.05, ##*p*<0.01, 0.3 mg/kg, \**p*<0.05, \*\**p*<0.01, \*\*\**p*<0.001, 1.0 mg/kg, versus vehicle (within-subject).

#### Cohort 2

Amphetamine reduced overall accuracy and increased incorrect responses at the highest dose (Fig 3d; Dose F_2,22_=21.52, *p*<0.001, vehicle versus 1.0 mg/kg *p* < 0.001, 0.3 mg/kg *p* = 0.465) but reduced omissions at all doses tested (Dose F_2,224_=7.36, *p*=0.004, vehicle versus 0.3mg/kg *p* = 0.031, 1.0 mg/kg *p* = 0.006). The number of correct trials was also reduced at the highest dose (Supplementary table S3; F_2,22_=16.41, *p*<0.001, vehicle versus 1.0mg/kg *p* < 0.001, 0.3 mg/kg *p=*0.970) but the total number of trials performed was not affected by treatment (Supplementary table S3; F_2,22_=2.98, *p*=0.072). Response latency was reduced by amphetamine treatment (Table 1; Dose F_2,22_ = 53.77, *p* < 0.001, vehicle versus 0.3 mg/kg *p* = 0.017, 1.0 mg/kg *p* < 0.001), and reduced the false-alarm latency at the highest dose (Table 1; Dose F_2,22_ = 16.83, *p* < 0.001, vehicle versus 1.0 mg/kg *p* < 0.001, 0.3 mg/kg *p* = 0.562). No other latency measures were affected (Table 1; Dose F_2,22_ < 3.18, *p* > 0.061). Further analysis showed that the highest dose increased responses to some distractor images; wheel and dots (Fig 3e; Image F_1.5,16.2_ = 46.09, *p* < 0.001, ε = 0.29, Dose F_2,22_ = 4.09, *p* = 0.031, Image*Dose F_5.5,60.4_ = 7.40, *p* < 0.001, ε = 0.55, vehicle versus wheel *p* = 0.017, dots *p* = 0.002). The highest dose increased accuracy when the target was presented first in the sequence (Fig 3f; Dose F_2,22_ = 21.63, *p* < 0.001, Position F_2.8,31.2_ = 83.65, *p* < 0.001, ε = 0.57, Position*Dose F_10,110_ = 8.31, *p* < 0.001, vehicle vs 1.0 mg/kg *p* = 0.019). Accuracy was reduced when waiting time for target presentation increased (target positions 2-5, 1.0 mg/kg *p* < 0.002, position 3 0.3 mg/kg *p* = 0.036), with no effect when the target was presented last (0.3mg/kg, 1.0 mg/kg *p* > 0.113).

### Atomoxetine Dose Response

Similar results were observed for both cohorts following atomoxetine treatment. Animals were overall more accurate but also made more omissions. The attention curves indicated that the improvement in accuracy was related to the presentation of the target during the later time points in the sequence whilst correct responses to the first image was reduced. In cohort 2, a specific improvement in discrimination between the target and false alarm image was observed.

#### Cohort 1

Atomoxetine treatment increased overall accuracy and decreased incorrect responses respectively (Fig 4a; Dose F_3,33_=5.19, *p*=0.005, vehicle versus 0.3 mg/kg *p* = 0.053, 1.0 mg/kg *p* < 0.001, 3.0 mg/kg *p* = 0.004). Correct trials and total trials were reduced at the highest dose (Supplementary table S3; F_3,33_=6.72, *p*=0.001, vehicle versus 3.0 mg/kg *p* = 0.031, and F_1.9,20.5_=17.29, *p*<0.001, ε = 0.62, vehicle versus 3.0 mg/kg *p* < 0.001 respectively). There was a dose-dependent increase in omissions (Fig 4a; Dose F_1.3, 13.8_ = 15.49, *p* = 0.001, ε = 0.42, vehicle versus 0.3 mg/kg *p* = 0.036, 1.0 mg/kg *p* < 0.001, 3.0 mg/kg *p* = 0.001), and response latency (Table 1; Dose F_3,33_ = 36.08, *p* < 0.001, vehicle versus 0.3 mg/kg *p* = 0.008, 1.0 mg/kg *p* < 0.001, 3.0 mg/kg *p* < 0.004). Collection and correct latency were also increased (Table 1; Dose F_3,33_ = 11.36, *p* < 0.001, vehicle versus 0.3 mg/kg *p* = 0.320, 1.0 mg/kg *p* = 0.001, 3.0 mg/kg *p* = 0.001, and Dose F_2.0,21.7_ = 6.66, *p* = 0.006, ε = 0.66, vehicle versus 0.3 mg/kg *p* = 0.135, 1.0 mg/kg *p* = 0.092, 3.0 mg/kg *p* = 0.010, respectively). No effects on incorrect latency were found (Dose F_2.1,22.8_=0.41, *p*=0.675). Analysis of image responses showed that atomoxetine reduced responses to some but not all the distractor images (Fig 4b; Image F_1.1,12.0_ = 93.16, *p* < 0.001, ε = 0.22, Dose F_3,33_ = 0.50, *p* = 0.682, Image*Dose F_15,165_ = 4.18, *p* < 0.001, flower 0.3 mg/kg, *p* = 0.022, 1.0 mg/kg *p* = 0.001, 3.0 mg/kg *p* = 0.024, dove 1.0 mg/kg *p* < 0.001, 3.0 mg/kg *p* = 0.003). Accuracy responses were reduced for the earliest presentation of the target image across all doses (Fig 4c; Dose F_3,33_ = 5.20, *p* = 0.005, Position F_3.4,36.8_ = 41.34, *p* < 0.001, ε = 0.67, Position*Dose F_15,165_ = 11.23, *p* < 0.001, 0.3 mg/kg *p* = 0.043, 1.0 mg/kg *p* = 0.001, 3.0 mg/kg *p* = 0.004), and for position 2 with the highest dose (p = 0.031). However atomoxetine increased accuracy when the target image was presented in the later target positions 3-6 (Position 3: 1.0 mg/kg *p* = 0.006, Position 4: 1.0 mg/kg *p* = 0.001, 3.0 mg/kg *p* = 0.021, Position 5: 0.3 mg/kg *p* = 0.029, 1.0 mg/kg *p* = 0.001, 3.0 mg/kg *p* < 0.001, Position 6: 1.0 mg/kg *p* < 0.001, 3.0 mg/kg *p* < 0.001).

**Figure 4.**
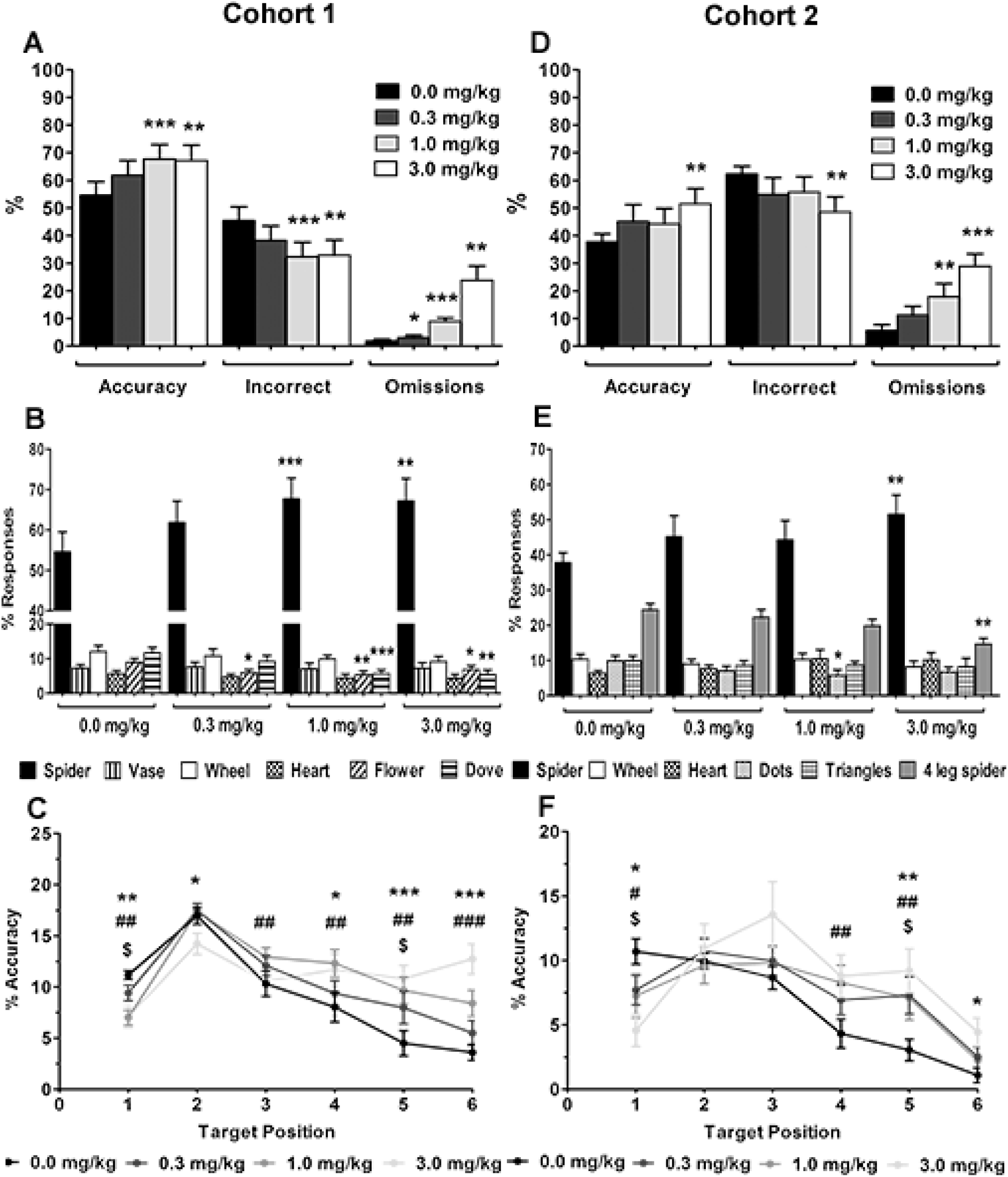
The effect of atomoxetine on performance in the rat-rapid serial visual presentation task (R-RSVP). Performance data for cohort 1 (a-c) and cohort 2 (d-f), response data for % accuracy, % incorrect, and % omissions for cohort 1 (a) and cohort 2 (d). Image responses for cohort 1 (b) and cohort 2 (e), spider is the target image. The sum of the responses to the distractor images (all images except spider) is equivalent to incorrect responses in (a). Attention curves showing accuracy per target sequence position for cohort 1 (c) and cohort 2 (f). Results are shown for the total population, mean ± SEM, *n* = 12 animals per cohort. Response data (a,b,d,e); \**p*<0.05, \*\**p*<0.01, \*\*\**p*<0.001, versus vehicle (within-subject). Accuracy per target sequence position (c,f); ^$^*p*<0.05, 0.3 mg/kg, #*p*<0.05, ##*p*<0.01, ###*p*<0.001, 1.0 mg/kg, \**p*<0.05, \*\**p*<0.01, \*\*\**p*<0.001, 3.0 mg/kg, versus vehicle (within-subject).

#### Cohort 2

Atomoxetine treatment increased overall accuracy and reduced incorrect responses at the highest dose (Fig 4d; Dose F_3,33_=5.13, *p*=0.005, 3.0 mg/kg *p* = 0.005). Omissions increased with atomoxetine treatment (Fig 4d; Dose F_3,33_ = 11.10, *p* < 0.001, 1.0 mg/kg *p* = 0.009, 3.0 mg/kg *p* < 0.001), as well as response latency and collection latency at all doses tested (Table 1; Response Latency: Dose F_1.8,20.0_ = 7.20, *p* = 0.005, ε = 0.61, 0.3 mg/kg *p* = 0.012, 1.0 mg/kg *p* < 0.001, 3.0 mg/kg *p* = 0.004, Collection Latency: Dose F_3,33_ = 3.37, *p* = 0.030, 0.3 mg/kg *p* = 0.020, 1.0 mg/kg *p* < 0.044, 3.0 mg/kg *p* = 0.024). Atomoxetine also reduced the number of correct and total trials performed (Supplementary table S3; F_3,33_=10.36, *p*<0.001, vehicle versus 3.0 mg/kg *p* < 0.001, and F_3,33_=35.73, *p*<0.001, vehicle versus 1.0 mg/kg *p* = 0.001, 3.0 mg/kg *p*<0.001 respectively). No other latency measures were affected (Table 1: Dose F_3,33_<1.90, *p*>0.149). Image response analysis showed that atomoxetine reduced responses to distractor images, with the main effect seen for the target vs false alarm although the lower dose also reduced responses to one of the other distractor images (Fig 4e; Image F_1.3,14.6_ = 35.95, *p* < 0.001, ε = 0.27, Dose F_3,33_ = 1.77, *p* = 0.172, Image*Dose F_15,165_ = 4.00, *p* < 0.001, 4-leg spider 3.0 mg/kg *p* = 0.002, dots 1.0 mg/kg *p* = 0.026). Atomoxetine reduced accuracy responses at the earliest target position for all doses versus vehicle treatment (Fig 4f; Dose F_3,33_ = 5.14, *p* = 0.005, Position F_5,55_ = 21.43, *p* < 0.001, Position*Dose F_15,165_ = 3.20, *p*<0.001, 0.3 mg/kg *p* = 0.034, 1.0 mg/kg *p* = 0.019, 3.0 mg/kg *p* = 0.019), but increased accuracy at later target positions 4-6 versus vehicle (Position 4: 1.0 mg/kg *p* = 0.004, Position 5: 0.3 mg/kg *p* = 0.012, 1.0 mg/kg *p* = 0.001, 3.0 mg/kg *p* = 0.030, Position 6: 3.0 mg/kg *p* = 0.030).

### Nicotine, Ketamine, Methylphenidate Dose Response

*Cohort 1 only:* One animal was excluded from the methylphenidate experiment due to a foot injury (n = 11). Nicotine reduced correct latency (Supplementary table S4; F_4, 44_= 5.73, *p* = 0.001, 0.3 mg/kg *p* = 0.003) and increased collection latency at the highest dose (Supplementary table S4; F_4, 44_= 4.22, *p* = 0.006, 0.3 mg/kg *p* = 0.043). No other performance variables were affected by nicotine treatment (Supplementary Fig S1, Supplementary Table S4; F_4,44_ < 1.25, *p* > 0.146). Ketamine increased omissions and reduced the total number of trials performed at the highest dose (Supplementary Fig S1, table S4; Dose F_3,33_=8.53, *p*<0.001, 10.0 mg/kg *p* = 0.003 and Dose F_3,33_=7.21, *p* = 0.001, 10.0 mg/kg *p* = 0.007, respectively). No other performance variables were affected by ketamine treatment (Dose F_3,33_<2.18, *p*>0.109). Methylphenidate had no effect on any performance variables (Supplementary Fig S1, Table S4; Dose F_3,33_<1.91, *p*>0.147).

## Discussion

These studies demonstrate that rats are able to learn and perform an RSVP-like attentional task involving a continuous sequence of images. Animals respond to a target image embedded in a randomized sequence of distractors demonstrating that this method can detect responses to go and no-go targets without having to use discrete trials or inter-image intervals. Accuracy levels were overall lower than in previous attentional tasks such as the 5CSRTT potentially making it easier to detect improvements in attention although it should be noted that only limited success was seen in terms of these pharmacological studies. Analysis of the attentional curves showed accuracy waned over time consistent with reduced ability to sustain attention or withhold responding. By having the concurrent measure of impulsivity across all images, we can also dissociate how drugs influence these two different variables and thus better understand whether the drug is influencing attentional processes or impulse control. The introduction of a false-alarm image gave a clear distinction between target (accuracy), false-alarm, and distractor responses, and suggests it may be possible to more clearly distinguish between specific impairments in discrimination or perceptual accuracy (responses to false-alarm image). We did not include a detailed analysis of the impact of different images and perceptual effects may influence the results although, by using a within-subject design for the drug studies, these are somewhat mitigated. It may also be that with further characterization of different image sets, we may be able to address this limitation and optimize the images and study design. Initial pharmacological investigations suggest that stimulant and non-stimulant treatments have different effects on animal’s performance in this task. Both amphetamine and atomoxetine treatment improved aspects of performance but with very different profiles in terms of the different performance measures recorded. Amphetamine’s effects were limited, and overall accuracy was reduced however, analysis of the attention curves revealed improvements in accuracy to the target when it was presented early in the sequence, possibly due to increased vigilance. In contrast, atomoxetine improved animal’s attention curves suggesting they were better able to sustain attention during the sequence presentation. Atomoxetine also specifically improved accuracy for the target versus the false alarm.

Training in the task took 42-60 sessions with the most challenging stage for the animals being the introduction of the no-go trials within the sequence. Modifications to the training procedure in future may further optimize this. The results from the baseline sessions confirmed that rats are able to distinguish a specific target image within a sequence of distractor images, including a false alarm image, presented in quick succession. Similar to other attentional tasks, we were able to measure overall accuracy (accuracy vs incorrect), omissions and both response and collection latencies. In this touchscreen RSVP task, including a false alarm image for cohort 2 meant errors of commission (incorrect responses) were more attributable to false-alarm responses than for other non-target (distractor) images. This enabled the dissociation between effects on discriminative accuracy or perception versus general impairments (responses to all distractor images arising from impulsive responding and/or omitted trials). Under normal conditions, animal’s overall probability of accurately responding to the target image decreased as a function of time which we suggest could provide a measure of sustained attention. These attentional data alongside measures of impulsive responding may also help differentiate between treatments which increase both impulsive responding and accuracy in rats as suggested by the results with amphetamine. Overall, we are able to measure similar outcomes to other attentional tasks but our initial studies suggest that this RSVP task may help dissociate between different aspects of attentional processing. Further investigations are needed but we suggest that integrating the findings from the different variables and analyses from this task may could potentially dissociate between, effects on sustained attention (attentional curve), discriminative or perceptual accuracy (target vs false alarm) and vigilance (omitted trials, response latency, attentional curves). These are in addition to similar measures of latencies and omissions used to understand effects on motivation and task engagement in the 5CSRTT [30]. A potential advantage of the RSVP task is that the animals cannot predict the presentation of stimuli thus reducing the influence of procedural learning and timing strategies^13,28^.

The level of accuracy shown by both cohorts should be sufficient to detect both improvements and impairments in attention with drugs that are known to alter attentional processing [15, 18]. This confers advantages over other tasks by allowing the detection of improvements without the need to either change task contingencies or use drug-induced impairments to reduce baseline performance [11, 31]. However, despite the reduced baseline accuracy seen in these studies is should be noted that, with the exception of atomoxetine, we did not observe improvements in accuracy in our acute pharmacology.

Amphetamine reduced accuracy and omissions in both cohorts and increased the speed of responding to the false-alarm image in cohort 2. Further analysis revealed that a reduction in response time was accompanied by a modest increase in target responses when presented at the earliest time point only. Performance in CPT are sensitive to the effects of stimulant drugs [32], with amphetamine improving performance and reaction times in normal humans, possibly through increasing vigilance and counteracting fatigue [9]. Translating amphetamine-mediated improvements in performance has been difficult in rodent attention tasks [15, 33, 34] although improvements in attention have been observed in human and mouse 5C-CPTs [35]. In this task amphetamine caused a modest increase in target responses at the earliest time point. However, overall there was no improvement in target vs false alarm discrimination and responses to all distractor images increased consistent with an increase in impulsive behavior [33, 36]. Amphetamine is known to increase responding in animals trained in operant tasks [15], through increases in dopamine release in areas such as the nucleus accumbens [37, 38]. The inability to wait for long periods in this task prevents the animal from responding to the target image in the later sequence positons, which likely contributed to the reduction in accuracy shown by amphetamine in this task.

The effects of atomoxetine were very different. Animals were more accurate overall and specifically showed greater ability to discriminate between the target and false alarm images. They were also better able to sustain their attention with more accurate responses made when the target was presented later in the sequence. However, they also made more omissions and latencies were increased suggesting there may be some more general effects on task engagement. Across both cohorts, atomoxetine increased accuracy of responding to the target image and improved accuracy in the attention curve analysis. In the second cohort, atomoxetine was found to also specifically increase accuracy for the target versus false alarm image. This improvement in the ability to distinguish between the target and false-alarm image infers that atomoxetine induced specific improvements in attention and the ability to inhibit distractor responding [39]. The lack of effect on the other distractor images also suggests that the attention curve effects were not related to a change in impulsive responding in this particular task. The increase in omitted trials is consistent with reported increases in omissions in the 5CSRTT[33]. Response latency was also increased indicating response suppression similar to that observed in a rat CPT using independent images [13] although the effects were not consistent. Based on the evidence from the attentional curves, atomoxetine appeared to improve the animal’s ability to wait before responding rather than affecting the ability to speed up recognition of and response to, target or non-target images. No changes in false-alarm latency or collection latency, further indicates that animals did not have slowed motor responses in general, [39]. Taken together these data suggest there is a tradeoff between speed and accuracy [40], rather than a change in motivation or lack of task engagement [41]. Animals responded less to the target image when presented in the 1^st^ or 2^nd^ sequence position but responded more when in the later positons. It is also interesting to observe how different these findings are from the touchscreen rCPT results [42] where decreased responding was observed across all measures.

In this study, oral doses of methylphenidate had no effect on any of the performance measures. The doses used were similar to those previously reported for the 5CSRTT and 5C-CPT [10, 18] but the use of an oral route of administration may have had an impact on the overall plasma levels achieved. Berridge et al., (2006) has previously suggested that the oral route of administration results in preferential effects on cortical versus sub-cortical dopamine potentially providing a more clinically relevant route of administration [43]. However, it should be noted that this study used animals which had already received other treatments which may have impacted on the sensitivity to this treatment. Methylphenidate administered to normal subjects typically reduces errors and reaction times in CPTs [44, 45]. Effects in rodent based tasks have mainly reported increases in impulsive responding at similar doses to that used here [18, 39, 46]. Modest improvements in attention in poor performing animals [18, 39] or differences in the pharmacokinetics of drug administration, may explain the lack of effect on attention shown here in animals performing optimally and with oral versus IP drug administration [18, 47]. Due to the small n number used in this study, analysis based on baseline performance levels was not carried out.

Ketamine had no specific effects on attention in this task but increased the level of omissions at higher doses, suggesting the animals became disrupted from performing the task. Ketamine’s lack of effect on attention is in-line with our previous study using a modified version of the 5-CSRTT that used an unpredictable stimulus presentation for a more attention demanding task [17, 48]. In normal human participants ketamine reduces accuracy trials, and increases omissions and incorrect trials in the AX-CPT [49, 50]. Ketamine-induced errors appear to specifically relate to responses to the target cue (‘X’) and inattention to the cue signal (‘A’) leading to increased responding to ‘B-X’ sequences versus other incorrect combinations that do not contain the target, i.e. ‘B-Y’ or ‘A-Y’ [49, 50]. This suggests that analyzing the type of error is important when assessing ketamine’s effect on performance.

Acute doses of nicotine had no effect on attention but did reduce correct latency in-line with previous reports using similar dose ranges and stimulus durations [51–53]. Nicotine-induced improvements in accuracy and response latencies have been reported previously in the 5CSRTT when task contingencies are changed unexpectedly resulting in impaired baseline performance [31, 54]. However, it is unclear as to how much non-attentional effects (response latencies) contribute to improvements in accuracy in this task. We also found an increase in collection latency at the highest dose which may reflect effects on the motivation for reward and food-rewarded behaviors [55]. This is difficult to compare to some key studies [31, 54] due to a lack of reporting on this parameter.

In drug-naïve humans effects of nicotine on attention are inconsistent across different task modalities with effects on commissions [56], omissions [7, 57], and reaction times [3, 56, 58], all being reported across variants of the CPT. In our task we found no specific effects on attention, or effects on omissions or distractor responses (commission errors). However, nicotine reduced correct latency and preserved target responding, which remains a consistent finding across CPTs [3, 11, 56]. The lack of effect of nicotine and ketamine may also have arisen as a result of animals undergoing multiple drug treatments and repeated testing in the task.

The different profile of effects of atomoxetine and amphetamine highlight possible differences between stimulant and non-stimulant effects on responding that may also impact on their clinical benefits in different types of ADHD [28, 59]. It has been suggested that the effects of these drugs involve similar catecholamine mechanisms within the prefrontal cortex [43, 60] however, our previous studies in noradrenergic lesioned animals suggested differences in their primary sites of action in the brain [61]. The findings from this rat RSVP task suggest that their effects may involve different mechanisms with amphetamine acting more on maintaining task and cue-elicited responding whereas atomoxetine improves attention by reducing the speed of responding and improving sustained attention. Further pharmacological studies and experiments involving disease models are needed to help extend knowledge of the validity of this task. These data provide initial support for this task in rats but there are limitations to the study. We only used a limited image data set and did not make a detailed assessment of the perceptual qualities of the different images and the studies here do suggest differences which could be better controlled. We also did not undertake the full pharmacological assessment in cohort 2 and pharmacokinetic issues with the methylphenidate study may have limited the plasma level achieved and hence the findings in cohort 1. Only male rats were tested and further work in females and in mice are necessary before the wider applicability of the task can be established. Finally, the study used a within-subject design and all animals received multiple drug treatments over the course of the experiment and carry over effects cannot be fully excluded although within experiment baseline data suggested the animals’ performance was stable.

## Acknowledgements

The authors would like to thank C. Hales for assistance with analysis using Matlab.

## Supplementary Materials

### Supplementary Methods

#### Training Procedure

Stage 1 of training involved a conditioned reinforcement schedule (CRF) to reinforce a touch response (nose poke or paw press) to any of the three blank (black) screens. The chamber light was switched off and only the magazine light illuminated for all training and testing. Each screen touch delivered a single reward pellet into the magazine (45 mg Noyes Precision Pellet, Sandown Scientific, UK) to a maximum of 100 pellets. Progression to the next stage occurred if >50 touch responses were made for 2 consecutive sessions.

Stage 2 involved responses to one screen only when a light grey background (LGB) image appeared. The LGB was randomly presented in equal numbers across each screen position (35 trials per screen location), and remained static until touched. Only one touch per trial was allowed, with new trials being initiated automatically. Touches on the blank (black) screens had no effect, only LGB responses were rewarded up to a maximum of 105 rewards (trials). Completion of stage 2 required >50 trials for 2 consecutive sessions. Stage 3 introduced punishment to non-LGB responses by a time-out period in which the house light was illuminated for 10 s. All image parameters the same as stage 2, criterion consisted of >50 trials for 2 consecutive days. For stage 4 animals were required to initiate each trial by nose poking in the magazine on the opposite wall of the chamber once illuminated. All other parameters were kept the same as the previous stage (criteria >50 trials, 2 consecutive days).

Stage 5 marked the start of the image sequence training in which the target image (‘spider’) was presented sequentially with two LGB images. The position of the target image within the sequence (1^st^, 2^nd^, or 3^rd^ position) was randomized and counterbalanced across the total number of trials (120 trials total). Therefore the waiting time for the presentation of the target image varied according to the position in the sequence. Once a trial was initiated each image in the sequence was presented for 3 s, with a total sequence time of 9 s. From this stage onwards, only the centre screen was used, with left/right screens remaining blank (black) and inactive. A response meant a single touch response on the target image and delivered a reward pellet. A single touch on either of the LGB images was classed as an incorrect response. An omitted trial meant that no responses had been made to any image presented during the sequence. Both incorrect and omitted trials were punished with a 10 s time-out and illumination of the house light. Possible trial outcomes, accuracy, incorrect, and omission, are illustrated in figure 1c. Only a single touch response per trial was allowed. To progress to the next stage animals had to perform >60% accuracy (chance performance = 33%), <30% omissions for 2 consecutive training sessions.

Stage 6 of training introduced distractor images into the image sequence. LGB images were replaced by 2 different picture images (Figure 1a, 1b), the total number of images remained at 3. The image sequence (including the distractor images) was randomized across the total number of trials (120 trials) to ensure animals could not predict the occurrence of the target from the presentation of the distractor images. All other parameters remained the same as the previous stage, criteria for progression was >60% accuracy, <20% omissions. For subsequent stages 7-9, additional distractor images were added until the image sequence contained 6 in total (target plus 5 distractors). A blank screen was presented before the image sequence to allow time for the animal to turn round and face the touchscreen once a trial had been initiated. An animal was considered trained once stage 9 was completed (>40% accuracy, <20% omissions, for two consecutive sessions). Chance performance using a 6-image sequence was approximately 17%.

### Additional training for cohort 2 (inclusion of a false alarm)

The false-alarm image (‘4-leg spider’) was modified from the target image, whereby 4 diagonally opposing legs were removed and a center circle of black pixels replaced by background pixels (figure 1b). Cohort 2 animals also underwent an additional training stage to reduce the image presentation time from 3 s to 2 s to further increase the attentional demands of the task as applied to other rodent attentional tasks. See video 1 for an example of baseline performance in the RSVP task (cohort 2).

---

**Supplementary Figure S1:**
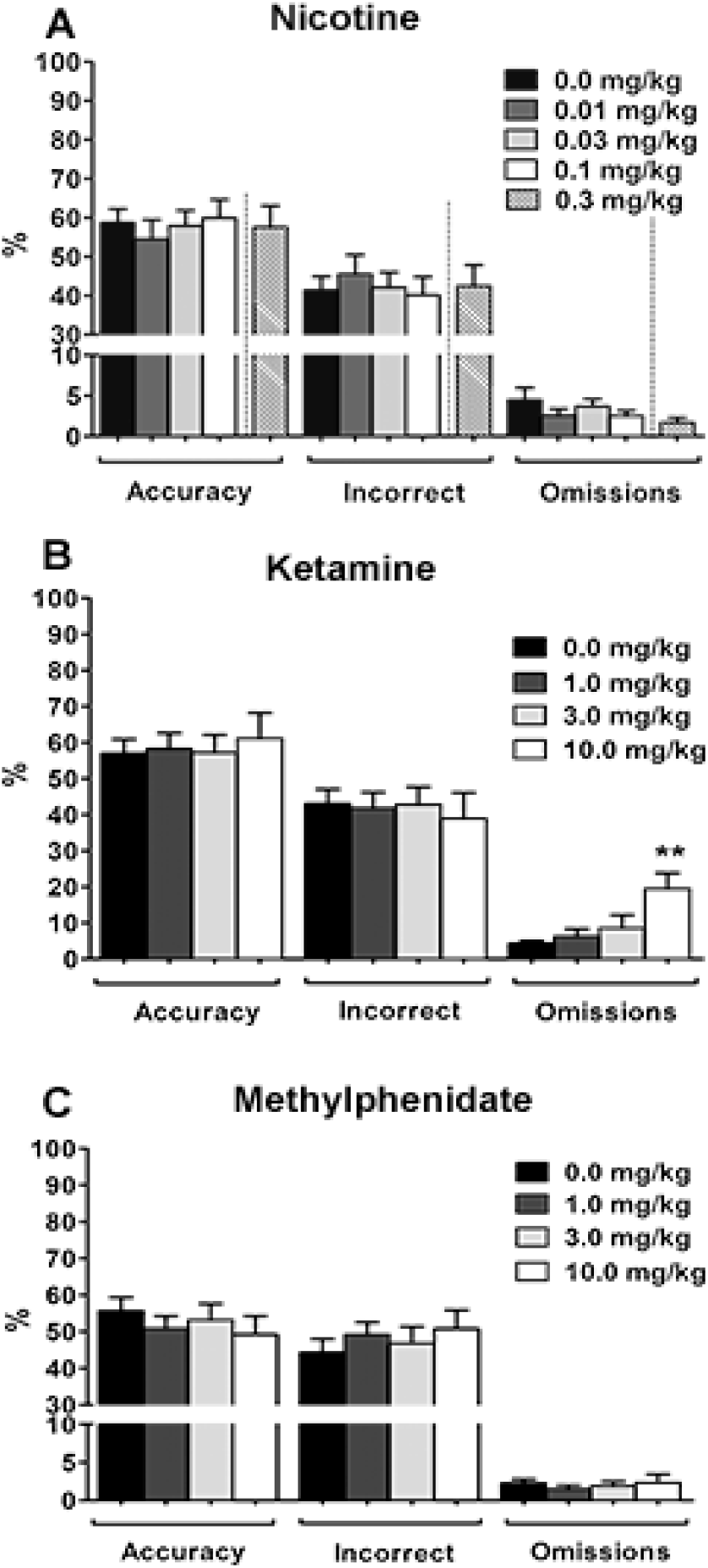
The effect of nicotine (a), ketamine (b), and methylphenidate (c), on performance variables % accuracy, % incorrect, % omission, in the rat-rapid serial visual presentation task (R-RSVP). Performance data is for cohort 1 only. Vertical lines indicate that the highest dose of nicotine (0.03 mg/kg) was administered to animals separately to the lower counterbalanced doses (a). Results are shown for the total population, mean ± SEM, *n* = 12 animals nicotine, ketamine, *n* = 11 methylphenidate, \*\**p*<0.01 versus vehicle (within-subject).

**Supplementary Table S1:** Training stages for rat-rapid serial visual presentation task.

| Stage | Description | Image Presentation | Criteria |
| --- | --- | --- | --- |
| 1. Touch Training: Conditioned Reinforcement | Images: 3 x Blank (black)<br>Screen touch = 1 reward pellet<br>100 rewards maximum | Continuous until touched | >50 touches<br>2 consecutive days |
| 2. Touch Training: One Image | Images: 1 x light grey background (LGB), 2 x blank (black)<br>LGB presented randomly in each screen position (left, right, centre)<br>LGB touch = 1 reward pellet<br>105 rewards maximum | Continuous until touched | >50 touches<br>2 consecutive days |
| 3. Introduction of Punishment | Same as previous stage except responses on blank (black) punished with 10 s time-out | Continuous until touched | >50 touches<br>2 consecutive days |
| 4. Trial Initiation | Same as previous stage with addition of trial initiation | Continuous until touched | >50 touches<br>2 consecutive days |
| 5. Sequence Training: Target Image | Images: 1 x target (spider), 2 x LGB<br>Random image order, presented in centre screen only<br>Target touch = 1 reward pellet ( trial)<br>LGB touch = 10 s time-out (in trial)<br>No touch = 10 s time-out (omitted trial)<br>120 trials maximum | 3 s / image | >60% accuracy<br><30% omissions<br>2 consecutive days |
| 6. Sequence Training: 3 images | Same as previous stage except LGB replaced with distractor images<br>Images: 1 x target (spider), 2 x distractor | 3 s / image | >60% accuracy<br><20% omission<br>2 consecutive days |
| 7. Sequence Training: 4 images | Same as previous stage with additional distractor image<br>Images: 1 x target (spider), 3 x distractor | 3 s / image | >60% accuracy<br><20% omission<br>2 consecutive days |
| 8. Sequence Training: 5 images | Same as previous stage with additional 'distractor' image<br>Images: 1 x target (spider), 4 x distractor | 3 s / image | >50% accuracy<br><20% omission<br>2 consecutive days |
| 9. Sequence Training: 6 images | Same as previous stage with additional 'distractor' image<br>Images: 1 x target (spider), 5 x distractor | 3 s / image | >40% accuracy<br><20% omission<br>2 consecutive days |
| 10. Reduced Presentation Time | Same as previous stage except reduced image presentation time | 2 s / image | >40% accuracy<br><20% omission<br>2 consecutive days |

**Supplementary Table S2:** Pre-drug data.

| Cohort | Total Trials | Correct Trials | Correct Latency (s) | Incorrect Latency (s) | False-Alarm Latency (s) | Response Latency (s) | Collection Latency (s) |
| --- | --- | --- | --- | --- | --- | --- | --- |
| 1 | 120 ± 0.1 | 69 ± 5.2 | 1.13 ± 0.05 | 1.45 ± 0.05 | - | 6.72 ± 0.32 | 1.52 ± 0.07 |
|  | 120 ± 0.0 | 69 ± 5.1 | 1.13 ± 0.05 | 1.45 ± 0.05 | - | 6.79 ± 0.30 | 1.50 ± 0.08 |
|  | 120 ± 0.0 | 71 ± 4.6 | 1.10 ± 0.04 | 1.50 ± 0.07 | - | 6.78 ± 0.35 | 1.51 ± 0.07 |
| 2 | 111 ± 2.7 | 37 ± 2.9 | 0.85 ± 0.09 | 0.98 ± 0.04 | 1.01 ± 0.04 | 4.03 ± 0.35 | 1.47 ± 0.07 |
|  | 114 ± 2.5 | 38 ± 3.9 | 0.87 ± 0.04 | 1.00 ± 0.03 | 0.97 ± 0.03 | 3.58 ± 0.27 | 1.46 ± 0.07 |
|  | 116 ± 1.8 | 39 ± 4.7 | 0.89 ± 0.05 | 0.97 ± 0.04 | 1.01 ± 0.03 | 3.75 ± 0.24 | 1.54 ± 0.10 |

Pre-drug latency data for cohort 1 (3 s image presentation) and cohort 2 (2 s image presentation) in the rat-rapid serial visual presentation task (R-RSVP). Only images used with cohort 2 contained a false-alarm image (4-leg spider), therefore false alarm latency for cohort 1 was not recorded. Results are shown for the total population, mean ± SEM, $n = 12$ animals per cohort.

**Supplementary Table S3:** Number of total and correct trials for amphetamine and atomoxetine.

| Cohort | Drug | Dose (mg/kg) | Total Trials | Correct Trials |
| --- | --- | --- | --- | --- |
| 1 | Amphetamine | 0.0 | 119 ± 0.7 | 68 ± 4.4 |
|  |  | 0.3 | 120 ± 0.0 | **59 ± 4.8\*\*** |
|  |  | 1.0 | 120 ± 0.0 | **38 ± 4.2\*\*\*** |
|  | Atomoxetine | 0.0 | 111 ± 5.4 | 59 ± 6.2 |
|  |  | 0.3 | 116 ± 4.1 | 69 ± 5.9 |
|  |  | 1.0 | 111 ± 3.9 | 67 ± 4.6 |
|  |  | 3.0 | **78 ± 8.4\*\*\*** | **42 ± 7.7\*** |
| 2 | Amphetamine | 0.0 | 111 ± 3.0 | 37 ± 3.2 |
|  |  | 0.3 | 117 ± 2.0 | 37 ± 3.1 |
|  |  | 1.0 | 118 ± 1.6 | **27 ± 1.8\*\*\*** |
|  | Atomoxetine | 0.0 | 111 ± 5.6 | 40 ± 3.8 |
|  |  | 0.3 | 106 ± 5.2 | 44 ± 7.1 |
|  |  | 1.0 | **80 ± 10.2\*\*** | 32 ± 6.2 |
|  |  | 3.0 | **46 ± 5.8\*\*\*** | **19 ± 3.9\*\*\*** |

Total number of trials performed and the number of responses for each drug dose tested in the rat-rapid serial visual presentation task (R-RSVP). Results are shown for cohort 1 (3 s image presentation) and cohort 2 (2 s image presentation), total population mean ± SEM, $n = 12$ animals per cohort, \* $p < 0.05$ , \*\* $p < 0.01$ , \*\*\* $p < 0.001$ , versus vehicle (within-subject).

**Supplementary Table S4:** Number of trials, responses, and latency data for nicotine, ketamine, and methylphenidate.

| Drug | Dose (mg/kg) | Total Trials | Correct Trials | Correct Latency (s) | Incorrect Latency (s) | Collection Latency (s) | Response Latency (s) |
| --- | --- | --- | --- | --- | --- | --- | --- |
| Nicotine | 0.0 | 114 ± 3.4 | 63 ± 4.0 | 0.97 ± 0.05 | 1.44 ± 0.05 | 1.64 ± 0.07 | 6.80 ± 0.31 |
|  | 0.01 | 118 ± 1.3 | 62 ± 5.3 | 0.96 ± 0.04 | 1.44 ± 0.06 | 1.57 ± 0.08 | 6.15 ± 0.40 |
|  | 0.03 | 118 ± 1.3 | 66 ± 4.7 | 0.96 ± 0.06 | 1.46 ± 0.03 | 1.64 ± 0.08 | 6.75 ± 0.22 |
|  | 0.1 | 120 ± 0.0 | 70 ± 5.4 | 0.95 ± 0.05 | 1.48 ± 0.04 | 1.59 ± 0.05 | 6.64 ± 0.35 |
|  | 0.3 | 120 ± 0.2 | 68 ± 6.2 | **0.83 ± 0.05\*\*** | 1.50 ± 0.04 | **1.77 ± 0.09\*** | 6.40 ± 0.38 |
| Ketamine | 0.0 | 107 ± 5.3 | 59 ± 5.7 | 0.96 ± 0.05 | 1.60 ± 0.05 | 1.72 ± 0.10 | 6.75 ± 0.30 |
|  | 1.0 | 102 ± 7.3 | 55 ± 5.8 | 1.10 ± 0.06 | 1.43 ± 0.05 | 1.62 ± 0.09 | 7.09 ± 0.42 |
|  | 3.0 | 105 ± 7.4 | 56 ± 7.0 | 1.10 ± 0.07 | 1.41 ± 0.05 | 1.67 ± 0.08 | 7.19 ± 0.55 |
|  | 10.0 | **77 ± 9.5\*\*** | 45 ± 8.7 | 0.97 ± 0.10 | 1.50 ± 0.09 | 1.66 ± 0.17 | 7.93 ± 0.44 |
| Methylphenidate | 0.0 | 114 ± 4.7 | 62 ± 5.3 | 0.92 ± 0.04 | 1.54 ± 0.04 | 1.60 ± 0.07 | 6.20 ± 0.29 |
|  | 1.0 | 119 ± 1.0 | 59 ± 4.0 | 0.95 ± 0.04 | 1.58 ± 0.05 | 1.50 ± 0.05 | 6.10 ± 0.18 |
|  | 3.0 | 120 ± 0.0 | 62 ± 5.2 | 0.92 ± 0.05 | 1.54 ± 0.04 | 1.54 ± 0.08 | 6.05 ± 0.31 |
|  | 10.0 | 116 ± 2.8 | 55 ± 5.5 | 0.98 ± 0.07 | 1.60 ± 0.05 | 1.56 ± 0.07 | 5.95 ± 0.50 |

The effect of nicotine, ketamine, and methylphenidate on total number of trials, the number of correct trials and latency measures, in the rat-rapid serial visual presentation task (R-RSVP) for cohort 1 (3 s image presentation). Results are shown for the total population, mean ± SEM, $n = 12$ animals nicotine and ketamine, $n = 11$ methylphenidate \* $p < 0.05$ , \*\* $p < 0.01$ , versus vehicle (within-subject).

